# Fractionated irradiation of murine salivary glands resulted in focal acinar cell atrophy, immune cell infiltration, fibrosis, and hyposalivation

**DOI:** 10.1101/2023.09.05.556313

**Authors:** Inga Solgård Juvkam, Olga Zlygosteva, Eirik Malinen, Nina Jeppesen Edin, Hilde Kanli Galtung, Tine Merete Søland

## Abstract

**Background:** Radiotherapy of head and neck cancer may cause detrimental late side effects such as fibrosis and hyposalivation. Our aim was to investigate late radiation-induced cellular and molecular changes of the salivary glands after fractionated irradiation to the head and neck in a murine model.

**Methods:** 12-week-old female C57BL/6J mice were irradiated with X-rays to a total dose of 66 Gy, given in 10 fractions over 5 days. The radiation field covered the oral cavity and major salivary glands. Salivary gland function was assessed by collecting saliva at baseline and at various time points after irradiation. The submandibular (SMG), sublingual (SLG), and parotid glands (PG) were dissected at day 105. Using different staining techniques, morphological, cellular, and molecular changes were investigated in the salivary glands.

**Results:** Saliva production was significantly reduced in irradiated compared to control mice at day 35, 80, and 105. We observed a significant decrease in total gland area and a significant increase in fibrotic area in irradiated compared to control SMG at day 105. Atrophy of acinar cells was observed in all irradiated SMG and SLG. Increased amount of chronic inflammatory cells, increased cell proliferation and altered expression of apoptotic markers were found in atrophic areas of irradiated glands.

**Conclusion:** Acinar and duct cells in irradiated salivary glands show increased cell proliferation and altered expression of apoptotic markers, proposing an attempt to overcome or withstand tissue damage caused by irradiation. This suggests a potential for regeneration of salivary glands after radiation therapy.

## Introduction

Head and neck cancer patients are often treated with radiotherapy which may cause severe late side effects such as tissue fibrosis and hyposalivation due to damages to normal tissue. Hyposalivation has detrimental effects on oral health causing loss of taste, difficulties in eating, speaking, and swallowing, increased incidence of caries and fungal infections, all with negative consequences for the patients’ quality of life [1, 2].

In humans and in rodents, the submandibular (SMG), sublingual (SLG) and parotid (PG), glands constitute the three pairs of major salivary glands. The salivary glands contain saliva-producing mucous and serous acinar cells, duct cells, myoepithelial cells, blood vessels and nerves that all can be affected by radiation [2, 3]. Despite having a slow tissue turnover rate, which usually corresponds to a late-responding tissue, the salivary glands also demonstrate an acute response to radiation. This acute response has been attributed to several factors such as dysfunction and apoptosis of acinar cells. The late response is thought to be due to chronic inflammation associated with glandular atrophy and repair by fibrosis resulting in tissue dysfunction, in addition to damages to the salivary gland stem cells [3-5]. Hyposalivation is thus considered both an acute and a late side effect after radiation [1]. However, the underlying biological mechanism of radiation-induced hyposalivation is not completely understood.

In a recent publication, we assessed several radiation-induced acute tissue responses such as oral mucositis, hyposalivation and skin dermatitis after fractionated X-ray irradiation in normal tissue of the head and neck in mice [6]. In the current work, our aim was to study late tissue responses in salivary glands after fractionated irradiation to the head and neck. We aimed at investigating radiation-induced morphological, cellular, and molecular changes of the salivary glands to further understand the underlying mechanisms behind hyposalivation.

## Materials and methods

### Animals

Nine-week-old C57BL/6J female mice were purchased from Janvier (Janvier Labs, France), kept in a 12-h light/12-h dark cycle under pathogen-free conditions and fed a standard commercial fodder with water given *ad libitum*. Standard housing with nesting material and refuge was provided. All experiments were approved by the Norwegian Food Safety Authority (ID 27931) and performed in accordance with directive 2010/63/EU on the protection of animals used for scientific purposes. At the onset of irradiation, animals were 12 weeks old.

### Irradiation Procedure

Using the preclinical model recently published [6], mice were irradiated with 10 fractions over 5 days (8 am and 4 pm) with a Faxitron Multirad225 irradiation system (Faxitron Bioptics, Tucson, AZ, USA) using following X-ray settings: 100 kV X-ray potential, 15 mA current and 2 mm Al filter with a dose rate of 0.75 Gy/min. Mice were randomly assigned to either sham treatment or 10 x 6.6 Gy (n = 10 for each treatment group). The radiation field was designed to include the oral cavity, pharynx, and major salivary glands while avoiding exposure to the eyes and brain. Mice were under anaesthesia during the irradiation procedure.

### Experimental Protocol

On day -7, baseline saliva collection was performed in all animals. On days 0 - 4, fractionated irradiation was given twice a day, as explained above. Saliva collection was then performed immediately after the end of fractionated irradiation (day 5), and at day 35, 80, and 105 during the follow-up period (Fig. 1). The saliva sampling procedure was performed as previously described [6, 7]. Saliva production was calculated as saliva volume (µl) per saliva collection time (15 minutes). The maximum follow-up period in this study was 105 days, in order to ensure full documentation of late tissue responses in the salivary glands, as fibrosis has previously been observed in mice at day 90 after single dose irradiation [5, 8]. At day 105, euthanasia was performed through overdose of anaesthetic (Pentobarbitol, Exagon® Vet) by intraperitoneal injection under terminal anaesthesia. The SMG, SLG, and PG were collected from each animal. All tissues were fixed in 10 % formalin for 24 hours before undergoing dehydration and embedding in paraffin.

**Figure 1.**
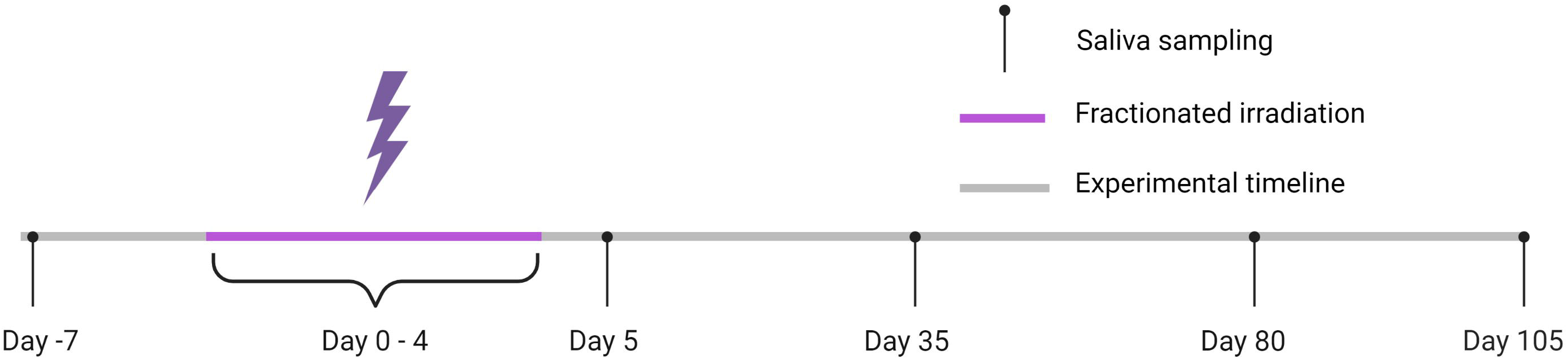
Timeline of the experimental protocol. Irradiation was performed twice a day for 5 days (days 0-4) in 10 fractions of 6.6 Gy. Saliva sampling occurred at baseline (day -7) and at several time points after irradiation (day 5, 35, 80, 105). Termination of the experiment was at day 105.

### Staining of tissues

For hematoxylin and eosin (HE), Alcian blue (AB), Masson trichrome (MT) and immunohistochemistry (IHC) staining, tissue sections of 4, 5, 6 and 4 µm, respectively were cut (Leica RM2155 microtome) and placed on microscopy glass slides (Superfrost Plus, Thermo Fisher) before deparaffination and hydration. For detailed staining procedures see *Supplementary materials*. After staining, all sections were dehydrated and mounted with xylene-based mounting medium (Pertex, Chemi-Teknik).

### Masson trichrome staining

MT was used to detect fibrosis as it stains fibrous tissue blue. MT staining was performed using a ready-to-use kit (Trichrome Stain kit, abcam) on a central section of each SMG and SLG. In the present study, there was a risk of incomplete dissection of PG due to its anatomical location. Since this could affect our results, PG was not included in the MT staining and the subsequent tissue analysis.

### Immunohistochemistry

Antigen retrieval was performed on deparaffinised sections of SMG and SLG, and the following primary antibodies were incubated over night at 4 LJ: rabbit anti-vimentin (clone EPR3776, Epitomics, dilution 1:500), rat anti-CD3 (clone CD3-12, MCA1477, Serotec, dilution 1:450), rabbit anti-CD45 (ab10558, abcam, dilution 1:1500), rat anti-F4/80 (clone BM8; eBioscience, dilution 1:400), rabbit anti-Bcl-2 (clone JF104-8; NOVUS, dilution 1:600), rabbit anti-BAX (clone 2F7I0; Invitrogen, dilution 1:300), and rabbit anti-Ki67 (clone SP6; Thermo Fisher Scientific, dilution 1:200). The secondary antibodies used were goat anti-rabbit IgG and rabbit anti-rat IgG. Cell nuclei were stained purple by Mayer’s hematoxylin.

### Analysis of salivary gland area and fibrotic area

For quantification of the total area of the salivary glands, overview images of MT-stained central sections of SMG and SLG were used and the total gland area was estimated by contouring the circumference of the gland using ImageJ. MT-stained sections were used to quantify the fibrotic area of the salivary glands, as this procedure is commonly used to analyse distribution of fibrotic tissue. With this method, cytoplasm and muscle are stained red, cell nuclei black, and collagen blue. The blue area can then be quantified as fibrotic area. Images of MT-stained sections of SMG and SLG were acquired using a Nikon DS-Ri1 camera with a CFI Plan Fluor 10x objective (NA 0.30). Several images (818x655 μm) of each gland were photographed to obtain a representative view of the entire gland. Using color thresholding in Image J, the fibrotic area in each image was measured relative to the total imaged area.

### Statistical analyses

Statistical analyses were performed in Prism 9 for Windows (Version 9.3.1, GraphPad Software, LLC). Saliva production was analysed using a mixed-effects analysis and Sidak’s multiple comparison test. Differences between irradiated and control groups with regards to the total gland area and fibrotic area were analysed using an unpaired t-test. A significance level of 0.05 was used for all comparisons.

## Results

### Decreased saliva production in irradiated mice

Saliva production was measured at baseline (day -7) and at different time points after onset of fractionated irradiation (day 5, 35, 80 and 105) (Fig. 1). The saliva production was significantly reduced in irradiated mice compared to controls at day 35 (p < 0.0001), day 80 (p = 0.0003) and day 105 (p = 0.03) (Fig. 2A).

**Figure 2.**
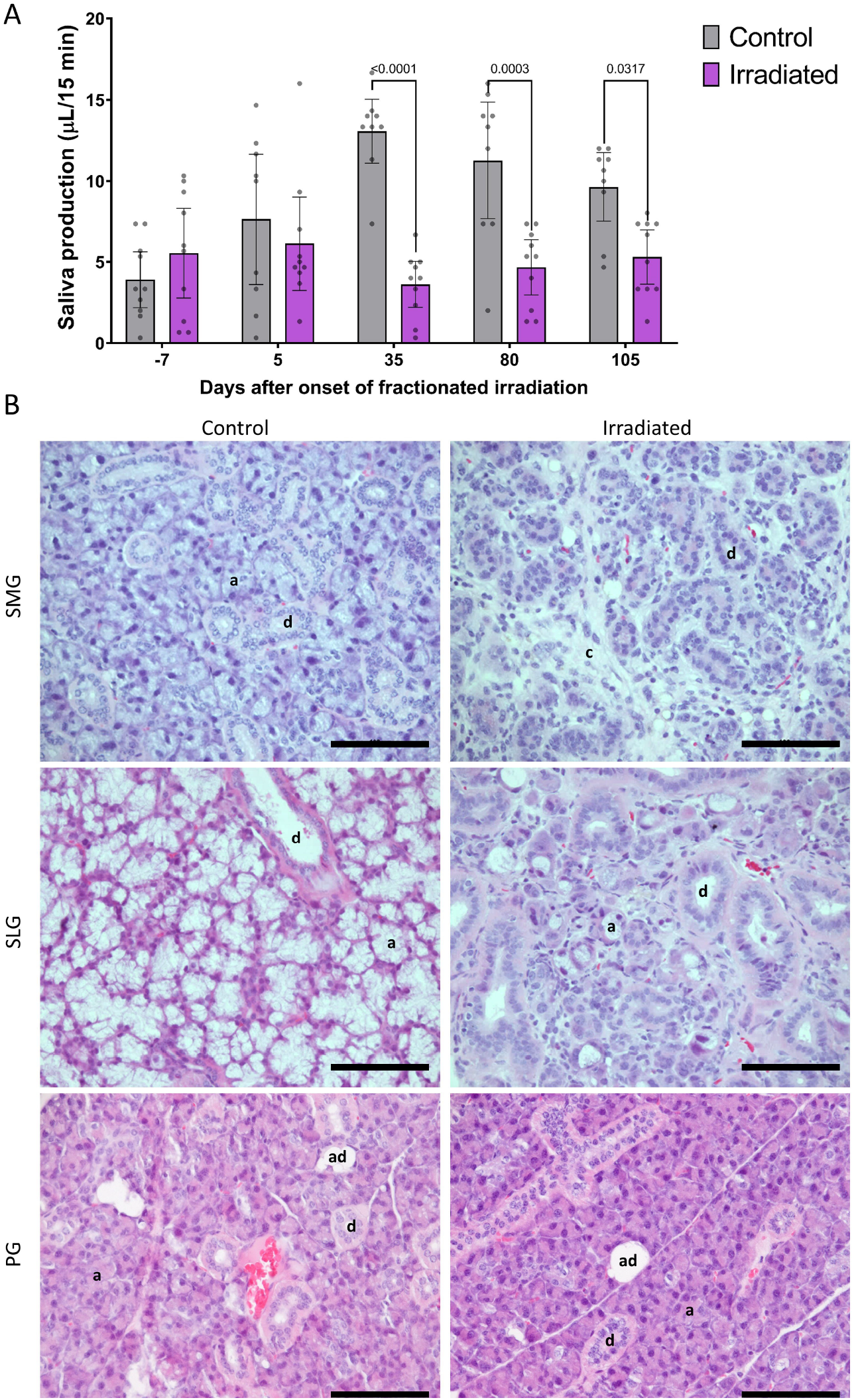
Decreased saliva production and acinar cell atrophy in SMG and SLG of irradiated mice. (A) Saliva production at baseline (day -7) and at various time points after onset of fractionated irradiation (day 5, 35, 80, 105). Dots represent data from individual animals. Data are presented as mean ± 95% CI. The p-value is shown between groups demonstrating statistically significant differences. (B) Representative images of hematoxylin and eosin (HE) stained sections 105 days after onset of fractionated irradiation in controls and irradiated mice. HE sections of irradiated and control submandibular (SMG), sublingual (SLG) and parotid (PG) glands, shows atrophy of acinar cells and their replacement by connective tissue in irradiated SMG and SLG but not PG. Scale bar is 100 µm. a = acinar cells, d = ducts, c = connective tissue, ad = adipocytes.

### Acinar cell atrophy in irradiated submandibular and sublingual glands

HE stained sections of the SMG and SLG showed loss of acinar cells in all irradiated mice at day 105 (Fig. 2B). No such changes were seen in the controls. Acinar cells were replaced by vimentin^+^ cells (Fig. 3A) and the tissue was stained blue by MT staining, both compatible with the presence of connective tissue. A reduced number of mucous acinar cells was demonstrated in irradiated versus control mice through AB staining (Fig. 3B). Interestingly, atrophy of acinar cells was observed only in one singular area of each irradiated gland and constituted approximately 40-70 % of the SMG and 20-70% of the SLG (Fig. 3C). HE stained sections of irradiated PG did not show atrophy of acinar cells.

**Figure 3.**
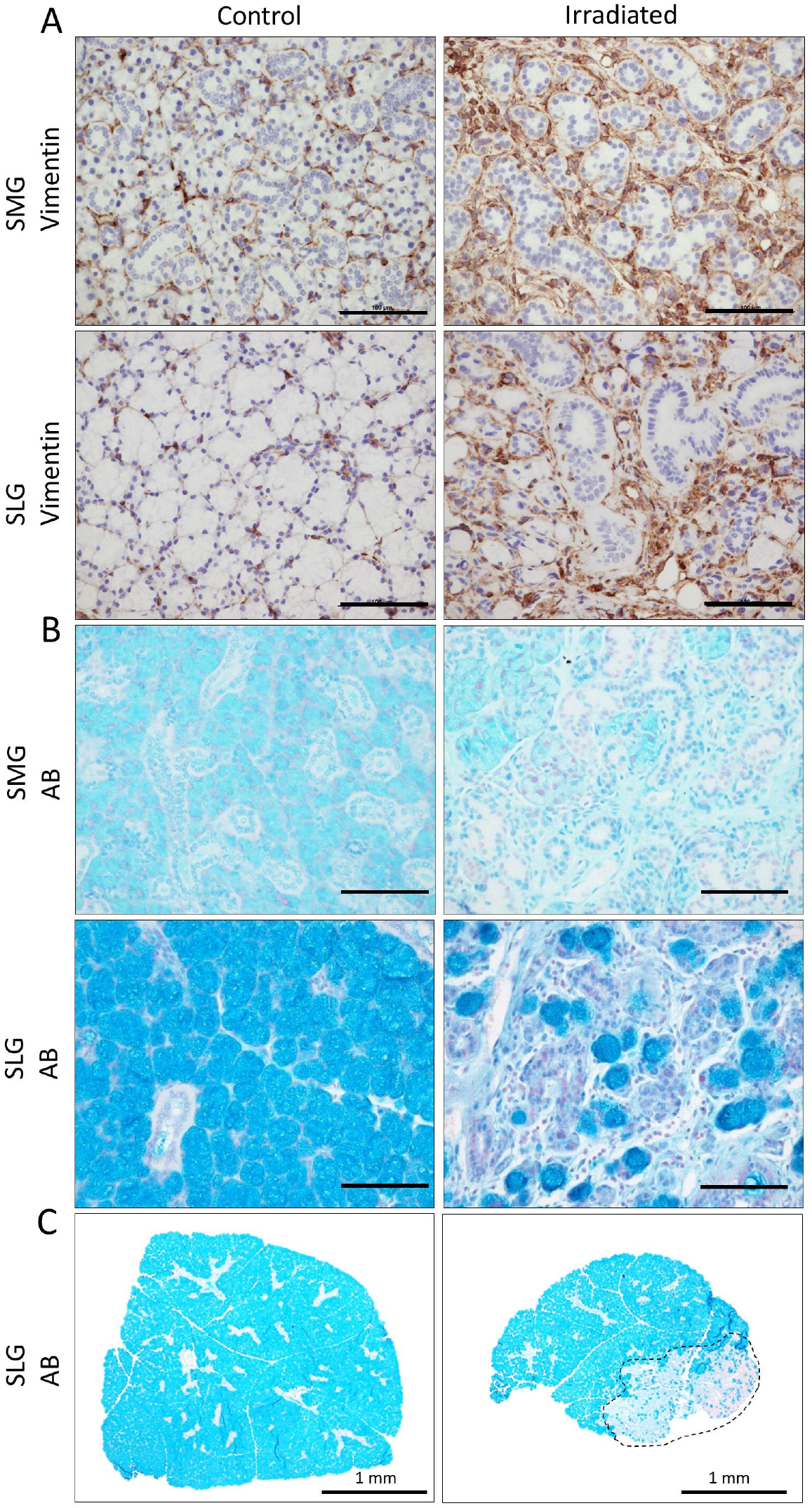
Increased vimentin expression and decreased number of mucous acinar cells in irradiated SMG and SLG. Representative images of the submandibular (SMG) and sublingual glands (SLG) at day 105 in controls and irradiated mice. (A) Sections labelled with an antibody against vimentin (brown), and cell nuclei stained with hematoxylin (purple). Increased number of vimentin^+^ cells is seen in the irradiated glands. Scale bar is 100 µm. (B) Mucous acinar cells are stained blue with alcian blue (AB), while cell nuclei are stained red with nuclear fast red. A reduced number of mucous acinar cells are seen in the irradiated sections. Scale bar is 100 µm. (C) Overview images of SLG (control and irradiated). The focal area (dotted line) of the irradiated SLG is stained less blue with AB staining both in comparison to the rest of the SLG tissue and the control.

### Decreased total salivary gland area and increased fibrotic area after irradiation

Based on MT-stained overview images, we quantified on average a 47 % smaller total gland area in irradiated SMG compared to controls at day 105 (p = 0.0001). Albeit not statistically significant, there was also a 21 % reduction in total SLG area in irradiated compared to control mice (p = 0.08) (Fig. 4A-B). From analysis of MT-stained sections, we found a significant increase of fibrotic area in SMG in irradiated compared to control mice (p = 0.0001) (Fig. 4C-D).

**Figure 4.**
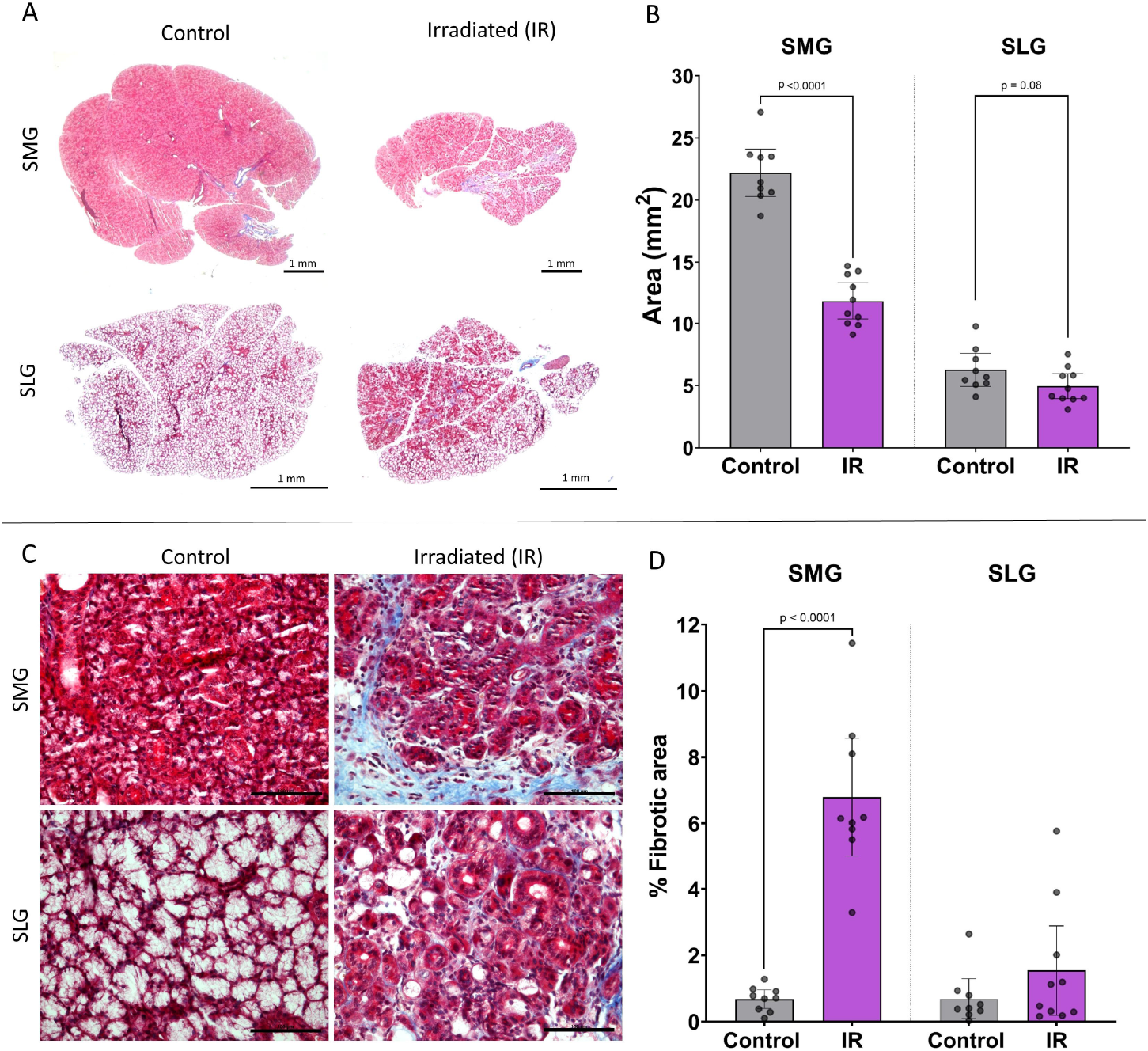
Decreased total gland area and increased fibrotic area in irradiated SMG and SLG. (A) Representative images of the entire gland in control and irradiated (IR) submandibular (SMG) and sublingual (SLG) glands used to quantify the total gland area. (B) Quantified total gland area in control and IR mice at day 105. Dots represents data from individual animals. Data are presented as mean ± 95% CI. (C) Representative images of Masson trichrome stained sections of SMG and SLG in control and IR mice. Collagen is stained blue and was quantified as fibrotic area. Scale bar is 100 µm. (D) Percentage of fibrotic area in SMG and SLG in control and IR mice at day 105. Dots represents data from individual animals. Data are presented as mean ± 95% CI. The p-value is shown between groups demonstrating statistically significant differences.

### Chronic inflammatory cells in irradiated salivary glands

Since fibrosis is known to be an end result of chronic inflammation [9, 10], we wished to study the inflammatory status of the irradiated SMG and SLG. We stained the sections with antibodies against CD3, CD45 and F4/80. CD3 is a marker for T-cells, CD45 is a general marker for leukocytes and F4/80 is a marker for macrophages in mice. In irradiated SMG and SLG, more CD3^+^ and CD45^+^ cells were present in the atrophic areas of the gland compared to the non-atrophic ones of the same gland and compared to non-irradiated control sections. This indicates and ongoing chronic inflammation in the atrophic areas of the irradiated glands. In addition, more F4/80^+^ cells were present in the atrophic area of irradiated SLG compared to the non-atrophic area of the same gland and compared to control sections. In contrast, F4/80^+^ cells were not observed in irradiated SMG, suggesting there are more macrophages present in the mucous SLG compared to the serous-mucous SMG (Fig. 5).

**Figure 5.**
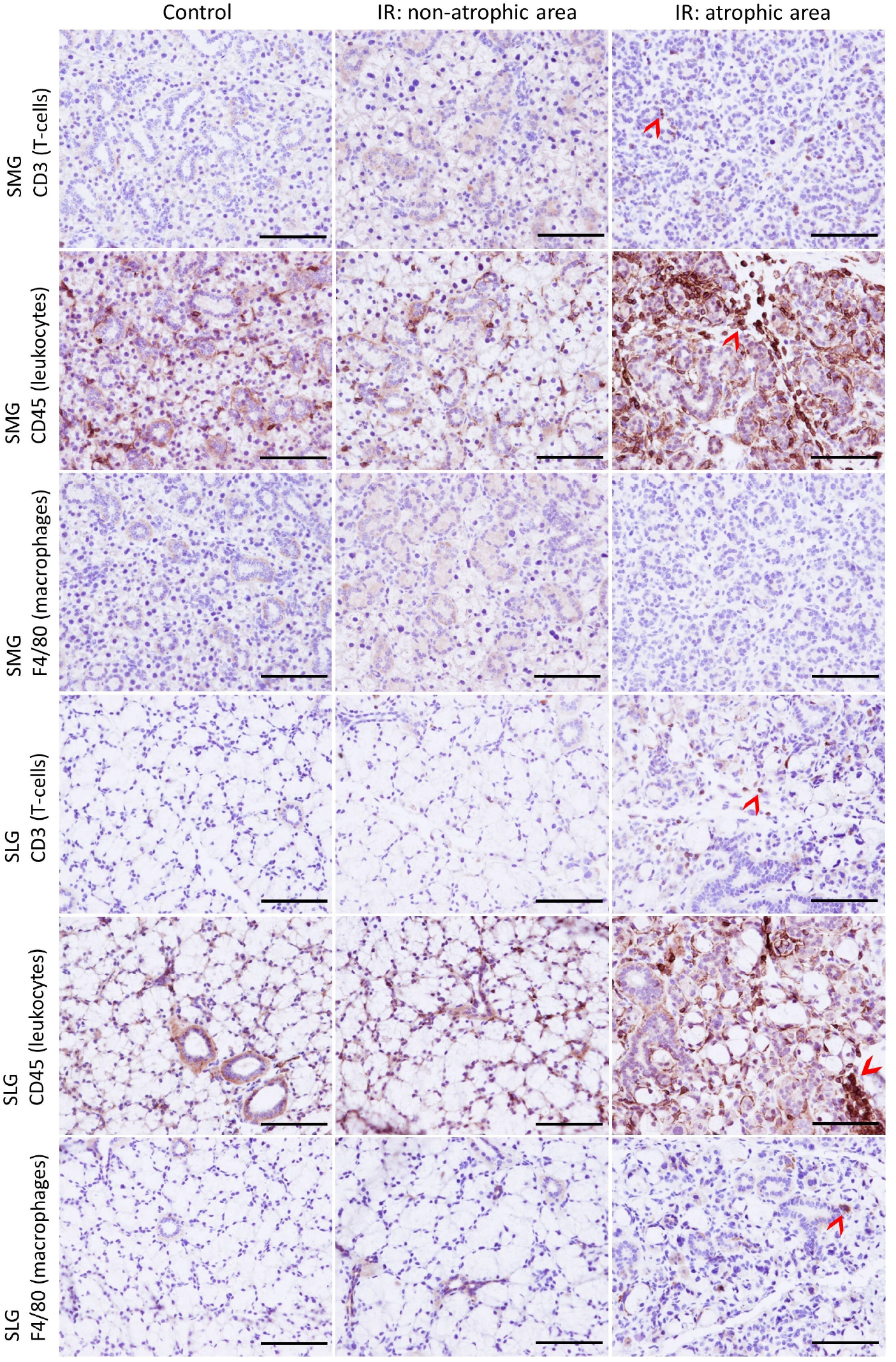
Chronic inflammatory cells in irradiated atrophic SMG and SLG. Representative images of atrophic and non-atrophic areas of irradiated (IR) and non-irradiated submandibular (SMG) and sublingual glands (SLG) immunohistochemically labelled with antibodies against CD3, CD45 and F4/80 (brown). Examples of positive cells are shown with red arrow heads. Cell nuclei were stained with hematoxylin (purple). An increased number of CD3^+^ and CD45^+^ cells were seen in the atrophic areas of IR SMG and SLG compared to the non-atrophic area in the same gland and control sections. More F4/80^+^ cells were present in atrophic SLG compared to atrophic SMG and controls. Scale bar is 100 µm.

### Altered expression of apoptotic markers and increased proliferation in irradiated salivary glands

High levels of apoptosis have been observed in fibrosis [11], as well as in atrophic salivary glands [12]. We therefore wanted to investigate apoptotic markers in the irradiated salivary glands. We stained the sections with antibodies against Bcl-2 and BAX, which are anti- and pro-apoptotic markers, respectively. An increased number of Bcl-2^+^ cells were observed in atrophic areas of irradiated SMG and SLG compared to the non-atrophic ones in the same gland and non-irradiated controls. Some of the Bcl-2^+^ cells were stellate spindle-shaped cells surrounding acinar cells (myoepithelial-like, as seen with α-SMA IHC staining in Suppl. Fig. 1) (Fig. 6), and some were coexpressing CD45 (Suppl. Fig. 2). We observed that the majority of Bcl-2^+^ cells in irradiated SMG and SLG were stromal cells. However, the remaining acinar and duct cells in irradiated SMG and SLG were Bcl-2^-^.

**Figure 6.**
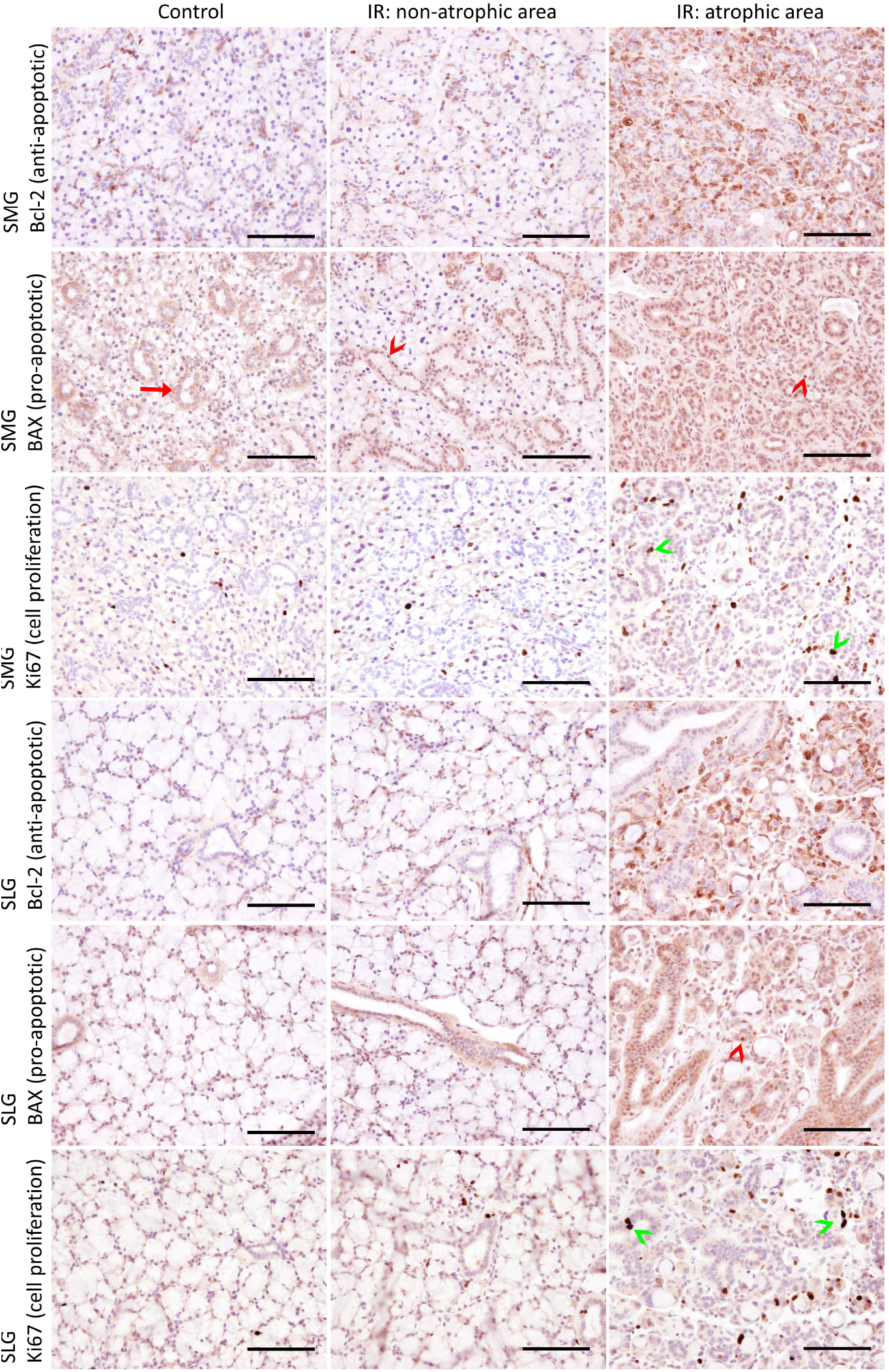
Increased proliferation and altered expression of apoptotic markers in irradiated atrophic SMG and SLG. Representative images of atrophic and non-atrophic areas of irradiated (IR) and non-irradiated submandibular (SMG) and sublingual glands (SLG) immunohistochemically labelled with antibodies against Bcl-2, BAX and Ki67 (brown). The cell nuclei were stained with hematoxylin (purple). More Bcl-2^+^ cells were observed in atrophic areas of IR SMG and SLG compared to non-atrophic areas of the same IR gland and controls. However, these Bcl-2^+^ cells were not acinar and ductal cells, but rather stromal cells correlating to the increased amount of stromal cells observed in the atrophic areas. BAX^+^ cells show more nuclear staining in non-atrophic and atrophic IR SMG (red arrow heads) compared to the cytoplasmic staining seen in controls (red arrow). We observed an increased amount of Ki67^+^ cells in atrophic parts of IR SMG and SLG. These Ki67^+^ cells were observed to be both acinar and duct cells (green arrow heads) as well as potential stromal cells. Scale bar is 100 µm.

Generally, an increased number of BAX^+^ cells were observed in atrophic areas of irradiated SMG and SLG compared to the non-atrophic areas of the same gland and non-irradiated controls. In irradiated non-atrophic and atrophic SMG we observed more nuclear location of BAX compared to the cytoplasmic staining seen in controls (Fig.6). Although BAX is considered a pro-apoptotic marker, nuclear BAX has been associated with non-apoptotic processes, for example by increasing cell proliferation of lung carcinoma epithelial cells [13]. To investigate cell proliferation in our study, we stained the sections with the cell proliferation marker Ki67. We observed Ki67^+^ acinar and duct cells in addition to an increased number of other Ki67^+^ cells in the irradiated atrophic areas of SMG and SLG (Fig. 6). Some of these other Ki67^+^ cells were most likely stromal cells or leukocytes as we observed an increase of vimentin^+^ cells (fibroblasts) and CD45^+^ cells (leukocytes) in the same area.

## Discussion

In the current study, we assessed late tissue responses after fractionated irradiation in terms of tissue fibrosis and hyposalivation in addition to investigating their underlying molecular mechanisms. Stimulated saliva production was significantly decreased in irradiated compared to control mice at day 35, 80 and 105 after onset of fractionated irradiation, indicating damages to the salivary glands. At day 105, we observed a significant decrease in total gland area and a significant increase in fibrotic area in irradiated compared to control SMG. Moreover, large areas of acinar cell atrophy accompanied by the presence of chronic inflammatory cells were observed in SMG and SLG. In contrast, atrophy of acinar cells and their replacement with fibrotic tissue were not observed in PG. Further molecular investigation revealed increased cell proliferation and altered expression of apoptotic markers, suggesting an attempt to overcome or withstand tissue damage caused by irradiation.

We have previously found histological changes in the PG, but not in the SMG and SLG at day 14 after fractionated irradiation [6]. In contrast, the current study demonstrated loss of acinar cells and their replacement by fibrotic tissue in SMG and SLG, but no loss of acinar cells in the PG, at day 105. A study using Wistar rats showed larger effects in the SMG than in the PG 240 days after fractionated irradiation [14]. In that work it was shown that after single dose irradiation, SMG and PG responded rather similarly, but their differences became evident after fractionated irradiation, especially at late time points after radiation. The investigators observed a larger reduction of gland weight and more loss of acinar cells in SMG than in PG, which is consistent with what we observed. Taken together, this may indicate that PG is more radioresistant compared to SMG and SLG, at least in mice and rats. However, due to the shielding of eyes and brain used in our study, it is possible that most but not the entire PG was included in the radiation field. This might be a limiting factor to our conclusion of differences in radioresistance between the three major salivary glands.

We found significantly reduced saliva production in irradiated compared to control mice after exposure to fractionated X-ray irradiation. Hyposalivation is a major problem both as an acute and a late side effect among head and neck cancer patients after irradiation and in patients with Sjögrens syndrome, a chronic inflammatory disease of salivary glands. In both conditions, the reduced saliva production can be due to loss of acinar cells and their replacement by fibrotic tissue [15, 16]. Loss of acinar cells and their replacement by fibrotic tissue has also been seen in rats and mice after single doses of X-ray irradiation [17-19]. In the present study we demonstrated loss of acinar cells and their replacement by connective tissue in SMG and SLG 105 days after onset of fractionated X-ray irradiation. Interestingly, the loss of acinar cells was only observed in certain areas of the irradiated SMG and SLG tissue. In these atrophic areas, we observed an increase of CD3^+^, F4/80^+^ and especially CD45^+^ cells, which indicates an ongoing chronic inflammation. An increase of CD45^+^ cells has also been observed 60 days after fractionated irradiation in the SMG of minipigs [20]. Due to the persistent nature of chronic inflammation [9, 10], it is expected that the atrophy of acinar cells and their replacement with fibrotic tissue would have continued after day 105, if the mice had not been terminated.

Since acinar cell atrophy was only observed in one singular area of each irradiated gland, we wanted to investigate whether the remaining acinar and duct cells in the same gland could have been protected from apoptosis through the anti-apoptotic protein Bcl-2 [21], and if acinar cells in atrophic areas were overexpressing the pro-apoptotic marker BAX [22]. Duct ligation is a method often used to study atrophy and regeneration of salivary glands and have several similarities with irradiation models [2]. After duct ligation, Bcl-2 has been found to be expressed by the remaining duct cells in atrophic SMG [12]. In contrast to their finding, the remaining acinar and duct cells in our study were Bcl-2^-^. This might suggest that they are not protected from apoptosis by Bcl-2. Moreover, we observed that many of the Bcl-2^+^ cells coexpressed CD45, a general marker for leukocytes, suggesting that the Bcl-2^+^ cells in our study are stromal and not parenchymal cells. Infiltrating mononuclear cells have also been seen to be Bcl-2^+^ in patients with Sjögrens syndrome [23]. Taken together, we found that the majority of Bcl-2^+^ cells in irradiated SMG and SLG were stromal cells and not acinar and duct cells. This may indicate that the remaining acinar and duct cells in irradiated glands are protected from apoptosis by a Bcl-2 independent pathway.

BAX is known to translocate from the cytosol to the mitochondrial membrane in response to apoptotic stimuli [22]. In healthy non-apoptotic cells, BAX is reported to remain inactive in the cytosol [24]. However, BAX has also been shown to translocate to the nucleus and function in relation to migration, differentiation and proliferation in non-apoptotic cells [13]. There the authors demonstrated that nuclear BAX increased proliferation in lung carcinoma epithelial cells. Cytosolic and nuclear location of BAX has also been observed in lung cancer cells [25, 26] and in human brain cells [27]. Thereby, it seems that the specific cellular location of BAX is important for its role and that cytosolic location represents an inactive form of BAX, mitochondrial BAX is involved in apoptosis, and that nuclear BAX can increase cell proliferation. In the present work, we observed increased nuclear location of BAX in atrophic areas of irradiated SMG and SLG, suggesting a cell proliferative and not a pro-apoptotic function of BAX. Indeed, we did observe Ki67^+^ acinar and duct cells in the same areas of irradiated SMG and SLG. Our observed increase of nuclear BAX support a role for this protein in cell proliferation in irradiated salivary glands.

In the current study, we have assessed late tissue responses such as chronic inflammation, tissue fibrosis, and hyposalivation in salivary glands after fractionated head and neck irradiation of mice. In particular, late tissue responses were only present in SMG and SLG suggesting less radioresistance of these glands compared to the PG. Furthermore, a protective anti-apoptotic mechanisms seems not to involve Bcl-2 in the remaining acinar and duct cells of irradiated SMG and SLG. Finally, we observed an increase of nuclear BAX and increased cell proliferation, which proposes a cell proliferative role and not only a pro-apoptotic function of BAX in irradiated salivary gland of mice. This is of great interest and suggests a potential for regeneration of salivary glands after radiation therapy.

## References

1. Siddiqui, F. and B. Movsas, Management of Radiation Toxicity in Head and Neck Cancers. Semin Radiat Oncol, 2017. 27(4): p. 340–349.

2. Chibly, A.M., et al., Salivary gland function, development, and regeneration. Physiol Rev, 2022. 102(3): p. 1495–1552.

3. Barazzuol, L., R.P. Coppes, and P. van Luijk, Prevention and treatment of radiotherapy-induced side effects. Mol Oncol, 2020. 14(7): p. 1538–1554.

4. Liu, Z., et al., Mechanism, Prevention, and Treatment of Radiation-Induced Salivary Gland Injury Related to Oxidative Stress. Antioxidants (Basel), 2021. 10(11).

5. Jasmer, K.J., et al., Radiation-Induced Salivary Gland Dysfunction: Mechanisms, Therapeutics and Future Directions. J Clin Med, 2020. 9(12).

6. Juvkam, I.S., et al., A preclinical model to investigate normal tissue damage following fractionated radiotherapy to the head and neck. J Radiat Res, 2022.

7. Bagavant, H., et al., A Method for the Measurement of Salivary Gland Function in Mice. J Vis Exp, 2018(131).

8. Urek, M.M., et al., Early and late effects of X-irradiation on submandibular gland: a morphological study in mice. Arch Med Res, 2005. 36(4): p. 339–43.

9. Wynn, T.A. and T.R. Ramalingam, Mechanisms of fibrosis: therapeutic translation for fibrotic disease. Nat Med, 2012. 18(7): p. 1028–40.

10. Wynn, T.A., Cellular and molecular mechanisms of fibrosis. J Pathol, 2008. 214(2): p. 199–210.

11. Johnson, A. and L.A. DiPietro, Apoptosis and angiogenesis: an evolving mechanism for fibrosis. FASEB J, 2013. 27(10): p. 3893–901.

12. Takahashi, S., et al., Cellular expression of Bcl-2 and Bax in atrophic submandibular glands of rats. Int J Exp Pathol, 2008. 89(5): p. 303–8.

13. Brayer, S., et al., The pro-apoptotic BAX protein influences cell growth and differentiation from the nucleus in healthy interphasic cells. Cell Cycle, 2017. 16(21): p. 2108–2118.

14. Coppes, R.P., A. Vissink, and A.W. Konings, Comparison of radiosensitivity of rat parotid and submandibular glands after different radiation schedules. Radiother Oncol, 2002. 63(3): p. 321–8.

15. Luitje, M.E., et al., Long-Term Maintenance of Acinar Cells in Human Submandibular Glands After Radiation Therapy. Int J Radiat Oncol Biol Phys, 2021. 109(4): p. 1028–1039.

16. Klein, A., et al., Acinar Atrophy, Fibrosis and Fatty Changes Are Significantly More Common than Sjogren’s Syndrome in Minor Salivary Gland Biopsies. Medicina (Kaunas), 2022. 58(2).

17. Kim, J.H., et al., Protective effects of alpha lipoic acid on radiation-induced salivary gland injury in rats. Oncotarget, 2016. 7(20): p. 29143–53.

18. Choi, J.S., et al., Enhanced tissue remodelling efficacy of adipose-derived mesenchymal stem cells using injectable matrices in radiation-damaged salivary gland model. J Tissue Eng Regen Med, 2018. 12(2): p. e695–e706.

19. Weng, P.L., et al., Limited Regeneration of Adult Salivary Glands after Severe Injury Involves Cellular Plasticity. Cell Rep, 2018. 24(6): p. 1464–1470 e3.

20. Lombaert, I.M.A., et al., CERE-120 Prevents Irradiation-Induced Hypofunction and Restores Immune Homeostasis in Porcine Salivary Glands. Mol Ther Methods Clin Dev, 2020. 18: p. 839–855.

21. Kroemer, G., The proto-oncogene Bcl-2 and its role in regulating apoptosis. Nat Med, 1997. 3(6): p. 614–20.

22. Murphy, K.M., et al., Bcl-2 inhibits Bax translocation from cytosol to mitochondria during drug-induced apoptosis of human tumor cells. Cell Death Differ, 2000. 7(1): p. 102–11.

23. Ohlsson, M., et al., CD40, CD154, Bax and Bcl-2 expression in Sjogren’s syndrome salivary glands: a putative anti-apoptotic role during its effector phases. Scand J Immunol, 2002. 56(6): p. 561–71.

24. Youle, R.J. and A. Strasser, The BCL-2 protein family: opposing activities that mediate cell death. Nat Rev Mol Cell Biol, 2008. 9(1): p. 47–59.

25. Salah-eldin, A., et al., Abnormal intracellular localization of Bax with a normal membrane anchor domain in human lung cancer cell lines. Jpn J Cancer Res, 2000. 91(12): p. 1269–77.

26. Bernal, C., et al., Regulatory Role of the RUNX2 Transcription Factor in Lung Cancer Apoptosis. Int J Cell Biol, 2022. 2022: p. 5198203.

27. Yao, Q., et al., Expression profile of the proapoptotic protein Bax in the human brain. Histochem Cell Biol, 2023. 159(2): p. 209–220.

