## Supplementary materials for "Fractionated irradiation of murine salivary glands resulted in focal acinar cell atrophy, immune cell infiltration, fibrosis, and hyposalivation"

### Supplementary material

*Hematoxylin and eosin staining*

By the use of progressive hematoxylin and eosin (HE) staining, the slides were immersed in Mayer's hematoxylin for 1 minute, followed by 2 minutes in 0.25 % hexamine and finally by 3 minutes in eosin. Between every staining step, the slides were washed in running tap water for 5 minutes.

*Alcian blue staining*

Alcian blue staining (AB) was used to detect mucous acinar cells as it stains acidic mucins blue. The slides were incubated with 3% acetic acid for 10 minutes at room temperature, washed in distilled water and incubated with Alcian blue (8GX, Sigma-Aldrich) for 30 minutes. Furthermore, the slides were washed in tap water and counterstained with nuclear fast red for 1 minute.

*Masson trichrome staining*

Masson trichrome (MT) staining was used to detect fibrosis as it stains fibrous tissue blue. MT staining was performed using a ready-to-use kit (Trichrome Stain kit, abcam) on a central section of each SMG and SLG. Briefly, the slides were immersed in preheated Bouin's fluid for 60 minutes at 60 °C. Thereafter, the slides were cooled, washed in tap water for 5 minutes, rinsed in distilled water and stained with Weigert's hematoxylin for 5 minutes. Subsequently, the slides were washed in tap water for 2 minutes and stained with Biebrich scarlet-acid fuchsin for 15 minutes. The slides were then rinsed in distilled water, incubated with phosphomolybdic-phosphotungstic acid for 15 minutes, dyed with aniline blue for 10 minutes, rinsed in distilled water and fixed with 1 % acetic acid for 5 minutes.

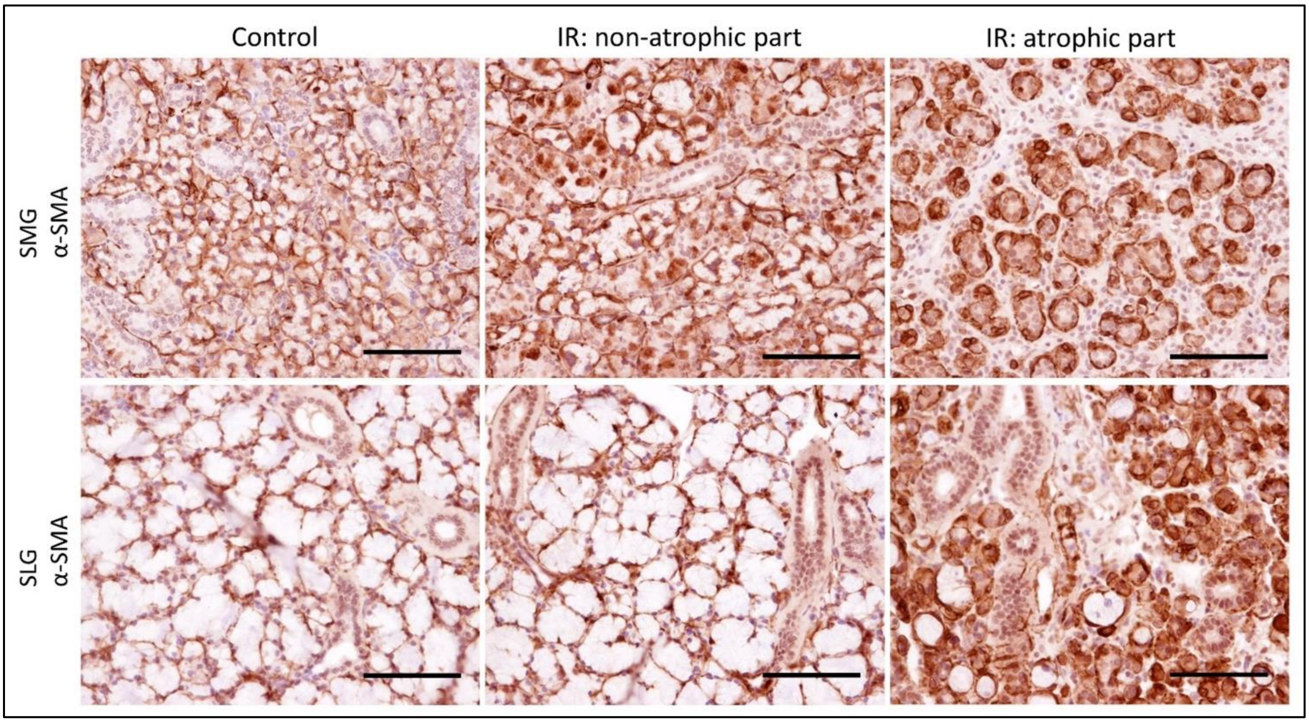


**Supplementary Figure 1.** Representative images of atrophic and non-atrophic parts of irradiated (IR) submandibular (SMG) and sublingual glands (SLG) and controls immunohistochemically labelled with antibodies against α-SMA (brown). α-SMA is used as a marker for myoepithelial cells that are stellate spindle-shaped cells surrounding acinar cells and occasionally intercalated ducts. Scale bar is 100 μm.

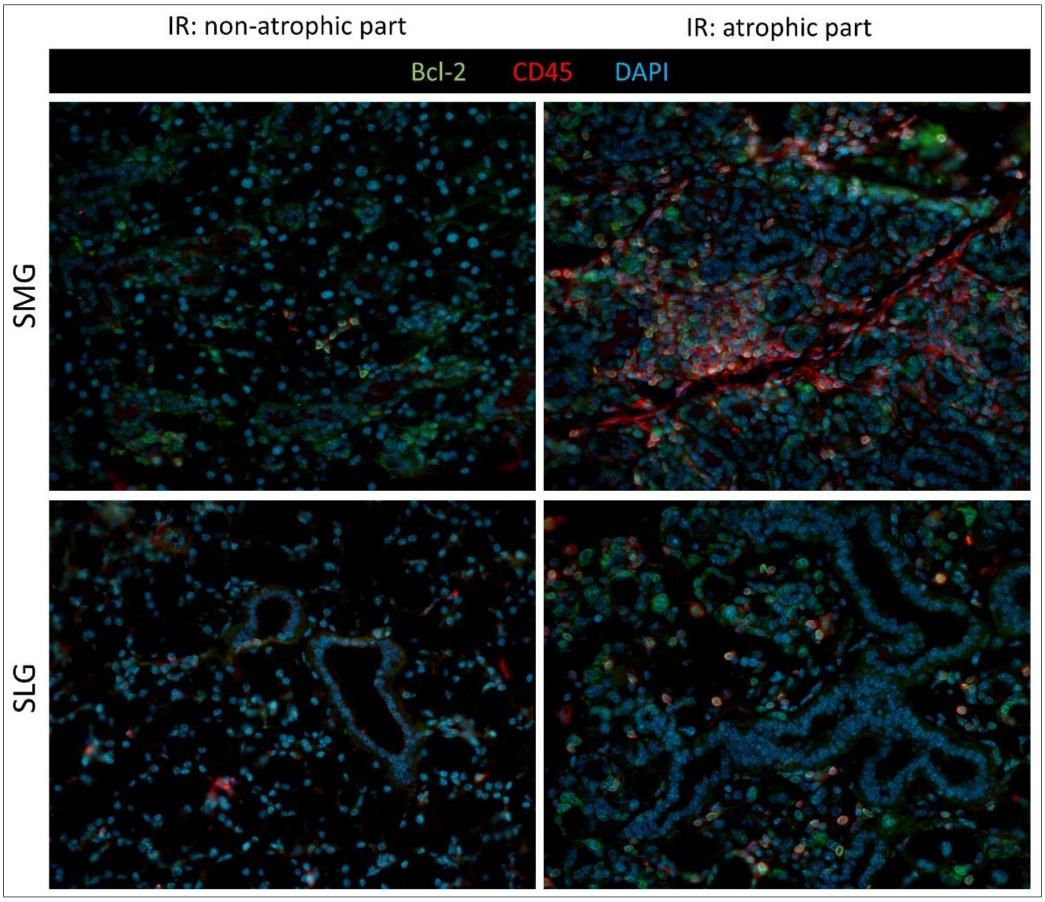

**Supplementary Figure 2. Many Bcl-2$^+$ cells coexpress CD45 in atrophic areas of irradiated SMG and SLG**

Representative images of atrophic and non-atrophic parts of irradiated (IR) submandibular (SMG) and sublingual glands (SLG) immunofluorescently labelled with antibodies against Bcl-2 (green) and CD45 (red). The sections were counterstained with DAPI to stain all cell nuclei (blue). Increased amount of Bcl-2$^+$ cells in atrophic parts of IR SMG and SLG, some of these also coexpress CD45. The images were taken with a 20x objective.
